# *Keetia korupensis sp. nov*. (Rubiaceae - Vanguerieae) a threatened lowland evergreen forest climber and the endemic plant species of Korup National Park, SW Region, Cameroon

**DOI:** 10.1101/2024.06.13.598689

**Authors:** Martin Cheek, Maguy Poundje, Laura Pearce

## Abstract

A review of the strictly endemic plant species of the Korup National Park, S.W. Region, Cameroon, is presented together with consideration of the importance of Korup for global plant conservation. Eighteen strict endemic species are known, 14 of which occur in lowland evergreen forest habitat (including *Keetia korupensis*) and of which 3 are among the 29 grove-forming monodominant detarioid legume species for which Korup is world famous. Four of Korup’s endemic species are rheophytes restricted to the Mana (Ndian) River rapids and falls which is probably the most species diverse site for shrubby rheophytes on the African continent. No endemics are yet known from Korup’s submontane forest, possibly because this is comparatively small in extent.

*Keetia korupensis* Cheek, a rainforest climber, is described and illustrated as strictly endemic, on current evidence, to the Korup National Park. The species differs from all other known species of the genus in its stipitate 2-seeded fruits which are twice as long as broad (vs non-stipitate and subisodiametric in most of the species) and for the 2 mm long stiff, skin-penetrating stem indumentum (vs non-injuring or soft in other long hairy species). *Keetia korupensis* is provisionally assessed using the IUCN standard as Vulnerable (VU D2).

## Introduction

*Keetia* E.P.Phillips (Rubiaceae, Vanguerieae) was segregated from *Canthium* Lam. by Bridson (1985, 1986). This genus of about 41 accepted species (Cheek & Onana 2024) is restricted to sub-Saharan Africa (excluding Madagascar and the Mascarene Islands), and extends from Senegal and Guinea in West Africa (Gosline *et al*. 2023a; 2023b) to Sudan (Darbyshire *et al*. 2015) in the North and East, and S. Africa in the South (Bridson 1986). *Keetia* differs from other African genera of Vanguerieae by its pyrenes with a fully or partly-defined lid-like area around a central crest, and endosperm with tanniniferous areas (Bridson 1986). *Keetia* species are usually climbers (very rarely shrubby) and occur mostly in forest habitats, sometimes in wooded grassland. In a phylogenetic analysis of the tribe based on morphology, nuclear ribosomal ITS and chloroplast *trnT-F* sequences, Lantz & Bremer (2004), found that based on a sample of four species, *Keetia* was monophyletic and sister to *Afrocanthium* (Bridson) Lantz & B. Bremer with strong support. Highest species diversity of *Keetia* is found in Cameroon and Tanzania, both of which have about 15 taxa (Onana 2011; POWO, continuously updated) and in Gabon, where only 10 species are currently recorded (Sosef *et al*. 2006) but around 25 are actually present, many of them undescribed (Lachenaud pers. comm. to Cheek, 2024). Several *Keetia* species are point endemics, or rare national endemics, and have been prioritized for conservation (e.g. Onana & Cheek 2011; Couch *et al*. 2019; Murphy *et al*. 2023; Darbyshire *et al*. 2023) and one threatened species, *Keetia susu* Cheek has a dedicated conservation action plan (Couch *et al*. 2022).

Bridson’s (1986) account of *Keetia* was preparatory to treatments of the Vanguerieae for the Flora of Tropical East Africa (Bridson & Verdcourt 1991) and Flora Zambesiaca (Bridson 1998). Pressed to deliver these, she stated that she could not dedicate sufficient time to a comprehensive revision of the species of *Keetia* outside these areas: “full revision of *Keetia* for the whole of Africa was not possible because the large number of taxa involved in West Africa, the Congo basin and Angola and the complex nature of some species would have caused an unacceptable delay in completion of some of the above Floras. […] A large number of new species remain to be described.” (Bridson 1986). Several of these new species were indicated by Bridson (1986), and other new species by her arrangement of specimens in folders that she annotated in the Kew Herbarium. One of these species was later taken up and published by Jongkind (2002) as *Keetia bridsoniae* Jongkind. In the same paper, Jongkind discovered and published *Keetia obovata* Jongkind based on material not seen by Bridson. Based mainly on new material, additional new species of *Keetia* have been published by Bridson & Robbrecht (1993), Bridson (1994), Cheek (2006), Lachenaud *et al*. (2017), Cheek *et al*. (2018a), Cheek & Bridson (2019), Cheek & Onana (2024)) and Cheek & Bissiengou (2024), and several other taxa that fit no other species, (e.g. Cheek *et al*. 2004; 2011) remain to be described.

In this paper we continue the project towards an updated taxonomic account of *Keetia* by describing a further new species, *K. korupensis* Cheek. This species was collected in 2008 as part of ongoing botanical surveys by RBG, Kew (K) and the IRAD-National Herbarium of Cameroon (YA) working towards completing the inventory of Cameroon’s plant species and to support conservation of its threatened species. The new species is highly distinctive for its skin-penetrating hispid hairs and elongated 2-seeded fruits, which are strongly stipitate and about twice as long as wide in side-view. Abundant material is available, but flowering material remains unknown.

New species to science are continually being described from Cameroon, mainly from the surviving species-diverse forests of the coastal plains and Cameroon Highlands. Among the most recent discoveries are new species of *Voacanga* (Apocynaceae, Lachenaud *et al*. 2024; Jongkind & Lachenaud 2022), *Drypetes* ( Putranjivaceae, Quintanar *et al*. 2023), *Didymochlaena* (Didymochlaenaceae, Shang & Zhang 2023), *Memecylon* (Melastomataceae, Stone *et al*. 2023), *Monanthotaxis* and *Uvariopsis* (Annonaceae, Cheek *et al*. 2023a; Gosline *et al*. 2022), *Impatiens* (Balsaminaceae, Janssens *et al*. 2022; Cheek *et al*. 2023b) and *Microcos* (Malvaceae-Grewioideae/Grewiaceae; Cheek *et al*. 2023c).

## Materials and Methods

Names of species and authors follow IPNI (continuously updated) and nomenclature follows Turland *et al*. (2018). Herbarium material was collected using the patrol method e.g. Cheek & Cable (1997). Identification and naming follow Cheek in Davies *et al*. (2023).

Herbarium specimens were examined with a Leica Wild M8 dissecting binocular microscope fitted with an eyepiece graticule measuring in units of 0.025 mm at maximum magnification. The drawing was made with the same equipment with a Leica 308700 camera lucida attachment. Pyrenes were characterized by boiling selected ripe fruits for several minutes in water until the flesh softened and could be removed. Finally, a toothbrush was used to clean the exposed pyrene surface to expose the surface sculpture. Specimens were inspected from the following herbaria: BM, FHO, K, P, YA. The format of the description follows those in other papers describing new species of *Keetia*, e.g. Cheek *et al*. (2018a). Terminology follows Beentje & Cheek (2003). All specimens cited have been seen. The conservation assessment follows the IUCN (2012) standard. Herbarium codes follow Index Herbariorum (Thiers, continuously updated).

### Taxonomic treatment

*Keetia korupensis* differs from all other known species of the genus in its stipitate 2-seeded fruits which are twice as long as broad (vs non-stipitate and subisodiametric in most of the species) and for the 2 mm long stiff, skin-penetrating stem indumentum (vs non-injuring or soft in other long hairy species).

It appears to have similarities (which may be superficial) to elements of the *Keetia hispida* (Benth.) Bridson complex rather than other species, in view of the hispid stem hairs and the broad caducous stipules. However, the two can be separated using the characters presented in Table 1.

**Table 1.** Characters distinguishing *Keetia hispida* and *K. korupensis sp*.*nov*. Data from specimens at K.

|  | *Keetia hispida* | *Keetia korupensis* sp. nov. |
| --- | --- | --- |
| Leaf texture | Thinly papery | Leathery |
| Fruit length: breadth ratio | c. 1:1 | c. 2:1 |
| Stipe present in fruit | Absent | Present, c. 5 x 2.5 mm |
| Shape of 2-seeded fruit | Subglobose to transversely ellipsoid | Clavate |
| Pedicel in fruit | Coiling | Remains straight |
| Nodes of primary stems | Hollow, ant-inhabited | Solid, not ant-inhabited |
| Stem indumentum | Hairs when hispid, < 2 mm long, not penetrating skin | Hairs dense, bristle-like, 2 mm long, penetrating skin |
| Pyrene base (side view) | Rounded | Acute |

### Keetia korupensis *Cheek* sp. nov

Type: Cameroon, S.W. Region, Korup National Park, North of P transect, N of P plot, 400 m east of edge of NW plot. Near *Tessmannia* grove, 5° 1’ 00”N, 8° 47’ 00”E. Primary forest with well-drained soil, fr. 22 Feb. 2008, *Pearce* 8 with van der Burgt, Corcoran, Elangwe (holotype K000593369; isotypes BR, EA, MO, P, SCA, YA). Fig. 1

**Fig. 1.**
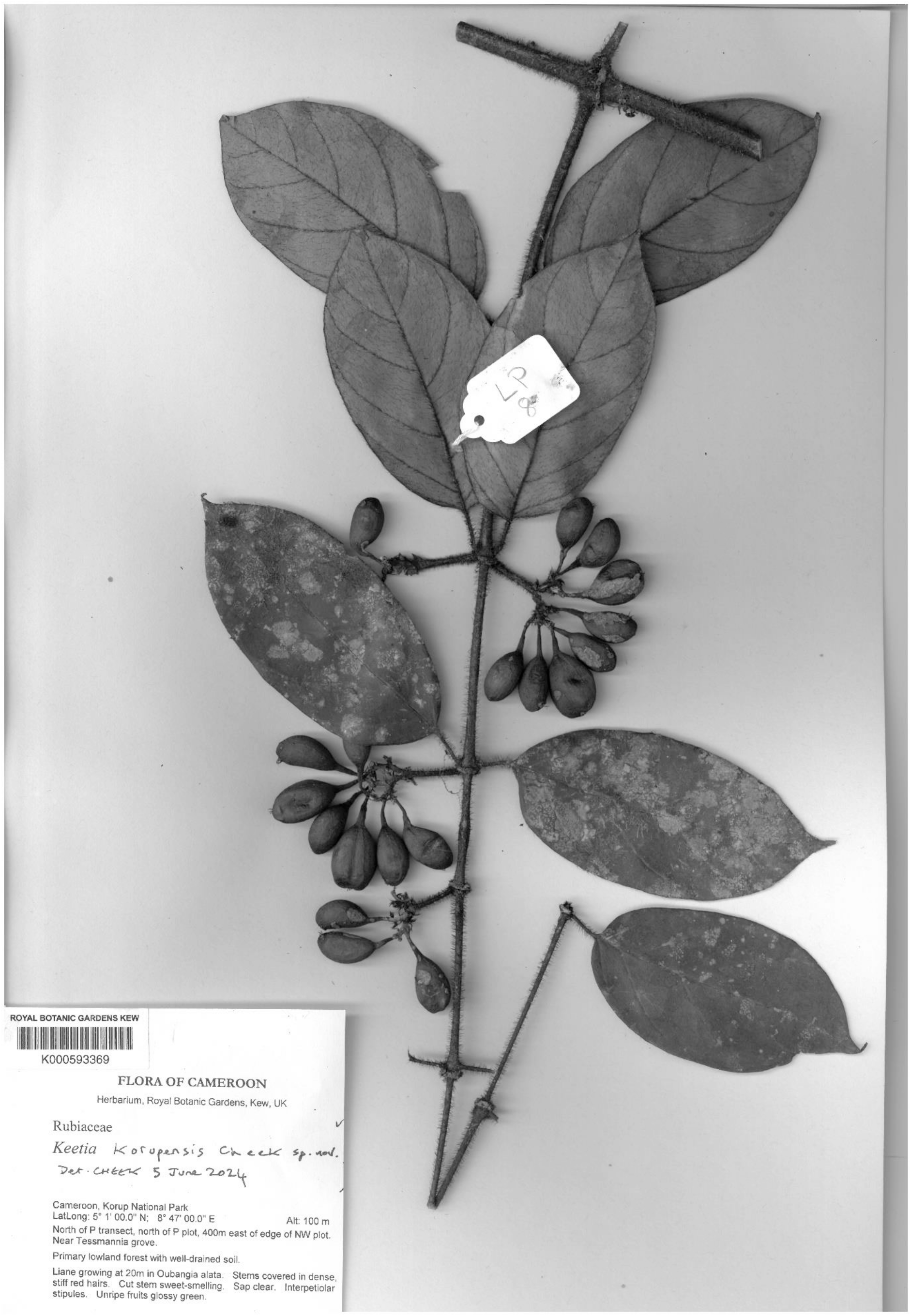
*Keetia korupensis sp. nov*. The holotype specimen, *Pearce* 8 (K, barcode K000593369).

*Liana* to 20 m tall. *Primary orthotropic stems* terete, internode length unknown, 3 – 6 mm diam., nodes thicker, 7 – 8 diam., the central 1 – 3 mm diam. with soft pith, sweet-smelling and producing clear liquid when cut (field notes), epidermis purple, lenticels not seen, densely persistent hispid hairy (piercing skin), hairs red (field notes), drying black, glossy, 1.5 – 2 mm long, c. 0.2 mm diam., apex acute (Fig. 1A-D).

*Secondary plagiotropic shoots* opposite, with (2 – )6 – 8 internodes, rarely branching further, as the primary stems but internodes lacking pith (3.6 – ) 5.5 – 7.7 x 0.18 – 0.35 cm, (Fig. 1A). *Leaves* of primary axis not seen; leaves of secondary shoots distichous, thickly coriaceous, drying dark matt grey green on upper surface, lower surface matt pale brown-white, elliptic to oblong, rarely lanceolate, (4.8 –) 9 – 10 (– 11) x (2 – )4.5 – 5.1( – 6.8) cm, more or less dimidiate, with one side up to 5 mm wider (from the midrib) than the other, acutely acuminate with acumen narrowly triangular c. 0.7 x 0.25 cm, twisting to one side, base obtuse or rounded to subtruncate, more or less asymmetric at base with one side more convex than the other, glabrous, margin slightly thickened and revolute, with a few long hairs (Fig. 1B); adaxial surface midrib slightly raised, with scattered hispid hairs as the stem, but mainly forward pointing, 0.4 – 1.2 mm long, secondary nerves slightly sunken, glabrous, blade with a few scattered hairs as midrib, at length glabrescent; abaxial surface with *domatia* and *bacterial nodules* absent (Fig. 1E); midrib strongly raised below, densely hispid hairy, hairs as stem, but shorter, 1– 1.25 mm long, secondary nerves (3 – )4 – 5 on each side of the midrib, raised and sparsely hispid below, arising from the midrib at 40 – 50°, straight, but near the margin curving upwards, parallel with and 2 – 3 mm from the margin, not uniting with the nerve above; tertiary and quaternary nerves inconspicuous, internerve areas with conspicuous dark brown, appressed, sparse hairs 0.9 – 1 mm long. *Petioles* black (dried material) terete to slightly plano-convex in transverse section, 0.8 – 1.2 x 0.1 – 0.12 cm, indumentum similar to stems. *Stipules* free, wider than the stem, soon reflexing from the base, drying thickly spongy, brown, fragile and soon breaking about 2 mm above the base and falling, not persistent apart from the shortly sheathing pale brown thickened spongy bases (fruiting stems), triangular, those of the primary stem not seen (apart from the persistent bases), those of the secondary stems (reconstructed) 7 – 10.5 x 5.5 – 6.6 cm, not keeled, slightly rounded at apex, abaxial (outer) surface (Fig. 1F) sparsely hairy with recurving thick black hairs 0.5 – 0.6 mm long; inner surface glabrous apart from a line of hairs at the base; hairs dense, black erect bristle-like c. 0.5 mm long with one or two colleters; colleters black, stout, cylindrical or gradually tapering to the apex, 0.5 – 1.25 x 0.2 – 0.25 mm.

*Inflorescences* (only known from fruit and old inflorescences) axillary on plagiotropic branches, in opposite axils or in only one axil, in 2 – 4 successive proximal nodes (not present in most proximal node); peduncles (in fruit) stout, patent or slightly ascending, 17 – 20 x 2 – 2.5 mm, hispid, hairs as in stem; main branches 2, short, 4 – 5 mm long, or so contracted as to be inconspicuous, the infructescence then appearing subcaptitate, subsequently 2 – 4-branched, further branches 3 – 4 mm long; bracts fallen, not seen; bracteoles subtending the base of the pedicels, foliose, flat, lanceolate or narrowly oblong, c. 3 x 0.5 mm, apex long acute, sparsely covered in hispid hairs c. 0.3 – 1 mm long. *Pedicel* terete 8 – 9 x 0.5 mm, glabrous, widening at apex into a stipe 2 – 4 x 2 mm *Calyx*-hypanthium obconical, 1.5 x 1.2 mm, moderately densely covered with stiff dark brown hairs, calyx tube shortly cylindrical, 0.2 mm long; teeth 5, linear to subspatulate, 0.9 – 1.1 x 0.2 mm, apices acute, outer surface with large hispid hairs c. 1 mm long (Fig. 1G), inner surface glabrous. *Corolla* not seen. *Disc* densely fine puberulent, domed, 1.5 – 2 mm diam., c. 0.5 mm long, the style base sunken (Fig. 1H). *Style* not seen.

*Fruits* green (field notes, possibly not fully mature), drying black-brown, fleshy, 1 – 2-seeded, 2-seeded fruits (1 in 5 of fruits) clavate oblong-elliptic (in side view), (1.7 – )1.9 – 2.8 x (1.0–) 1.3 – 1.4 x 0.9 cm, very slightly didymous, with a shallow, barely discernible groove running from stipe to calyx, apex truncate, the calyx 3 mm wide in a small central excavation, calyx tube to 0.5 mm long; teeth persistent, linear, 1 – 2 mm long; stipe subcylindrical, widest at apex, 3 – 5 x 1 mm diam. at junction with pedicel, to 2.5 mm diam. at apex, glabrous (Fig. 1J & K); 1-seeded fruits as 2-seeded but slightly curved, oblong (1.6 – )1.8 – 2.2 x (0.8 –) 0.9 – 1.05(– 1.15) x 0.9 cm, calyx subapical.

*Pyrenes* pale brown, crustaceous, thinly woody, smooth, unsculptured, those of 2-seeded fruits narrowly obovoid with flattened ventral side, (16 –)18 – 19 x 7.5 x 8 mm; pyrene wall 0.3 – 0.6 mm thick; lid completely apical, cap-like, orbicular, c. 4 mm diam., 2 mm long, with a low, broad, inconspicuous central crest; dehiscing (during preparation, possible artefact) along the midline of the crest and the dorsal (abaxial) edge of the cap (Fig. 1L); point of attachment immediately below the cap, cap held across top of the pyrene (at right angles to the ventral face) (in 2-seeded fruits) or angled c. 20 degrees towards the ventral face (in 1-seeded fruits). *Seed* ellipsoid, c. 10 x 5.5 – 6 x 3 mm (possibly immature), surface black to dark brown; embryo (transverse section) central, surrounded by dense tanniniferous rays (Fig. 1M).

## RECOGNITION

Differs from all known species of the genus in having 2 mm long, skin penetrating hispid hairs on the stem (vs usually <1.5 mm, not skin penetrating), fruits clavate, length: breadth ratio c.2: 1, including a pronounced stipe to 5 x 2.5 mm (vs subglobose or ellipsoid, non-stipitate); pyrenes which are narrowly obovoid, length: breadth ratio 2:1 (vs subglobose, <1.5:1) and with pyrene bases acute in side view, not rounded; otherwise similar to *K. hispida* but lacking ant-filled nodal cavities. See Table 1 for additional diagnostic characters.

## DISTRIBUTION

Korup National Park, S.W. Region, Cameroon

## SPECIMEN EXAMINED. CAMEROON

S.W. Region, Korup National Park, North of P transect, N of P plot, 400 m east of edge of NW plot, fr. 22 Feb. 2008, *Pearce* 8 with van der Burgt, Corcoran, Elangwe (holotype K000593369; isotypes BR, EA, MO, P, SCA, YA).

## HABITAT & ECOLOGY

Lowland evergreen forest with *Oubanguia alata* Baker f. (Lecythidaceae-Scytopetaloideae, fide *Pearce* 8) c. 100 m alt.

## CONSERVATION STATUS

*Keetia korupensis* is known from a single site in the Korup National Park, where threats are currently thought to be low and forest is largely intact apart from in the vicinity of current or former enclaved villages. However, since December 2016 the people of NorthWest and SouthWest Regions have sought to secede from the rest of Cameroon and the consequent armed conflict has resulted in the deplacement of much of the population into Nigeria and neighbouring Regions. Some communities have moved into protected forest areas which poses a threat to biodiversity conservation, as in Lebialem, also in SW Region (Tabi *et al*. 2020). It is possible that this is the case in Korup, which would pose a threat to this species. *Keetia korupensis* appears to be extremely rare since only a single collection, with only a single individual is known for certainty. This is despite numerous large scale inventory surveys for plants in Korup between 1980 and 2016, including multiple intensively sampled plot areas both large and small over the mid and southern parts of the National Park (Thomas *et al*. 2003; van der Burgt *et al*. 2021; Cheek & Cable 1998). Such survey efforts themselves have posed a level of threat, albeit at low intensity, since in conducting them, temporary camps, and paths, can result in the low level cutting of plants. The area of occurrence (AOO) is estimated as 4 km^2^ using the IUCN required grid cells of that size, although in reality it is known from a much smaller area. The extent of occurrence (EOO) of this taxon is inferred as 5 km^2^ since according to IUCN rules, this must be larger than the AOO.

Therefore, *Keetia korupensis* is provisionally assessed as Vulnerable (VU D2). It is to be hoped that the species will be found at further sites in the Korup or in neighbouring areas, however the prospect of this seems low since numerous intensive botanical surveys have been conducted in areas to the N and E of Korup without encountering this species (Cable & Cheek 1998; Chapman & Chapman 2001; Cheek *et al*. 1992;1996; 2000; 2004; 2006; 2010; 2011; Harvey *et al*. 2004; 2010; Maisels *et al*. 2000).

Seventeen other species are also currently globally unique to the Korup, e.g. (see discussion, below).

## ETYMOLOGY

The species is named for Cameroon’s Korup National Park where it was discovered and to which it appears unique globally

## Notes

*Keetia korupensis* is perhaps most likely to be mistaken for *K. hispida* s.l. especially the variants previously treated as *Canthium setosa* Hiern, due to the long patent stem hairs, and similar size and shape of leaf blades. This last taxon is relatively frequent in SW Region, Cameroon. However, the two taxa can be easily separated as shown in Table 1. Less frequent in Cameroon is *K. molundensis* (K. Krause) Bridson, which shares with *K. korupensis* the unusual large, broad, flat stipules lacking the subulate awn seen in most species of the genus. However, this species has subglobose fruits, domatia that extend along the secondary nerves (vs elongate, stipitate fruits; no domatia), and stem hairs that are soft, and which dry red (vs stiff, skin puncturing and drying black).

## Discussion

### Korup National Park: of global importance for plant conservation

Korup National Park (1873 km^2^) has the highest number of nationally endemic and range-restricted (extent of occurrence <10,000 km^2^) plant species (85 taxa) recorded in Cameroon, after Ngovayang (88 taxa) (Murphy *et al*. 2023). 147 of these Korup taxa are globally threatened, in the categories Critically Endangered (17 species), Endangered (45 species) and Vulnerable (85 species) and Korup is designated as an Important Plant Area (Murphy *et al*. 2023). With the publication of *Keetia korupensis*, the number of globally endemic (unique to Korup) species rises to 18 (see Table 2).

**Table 2.**
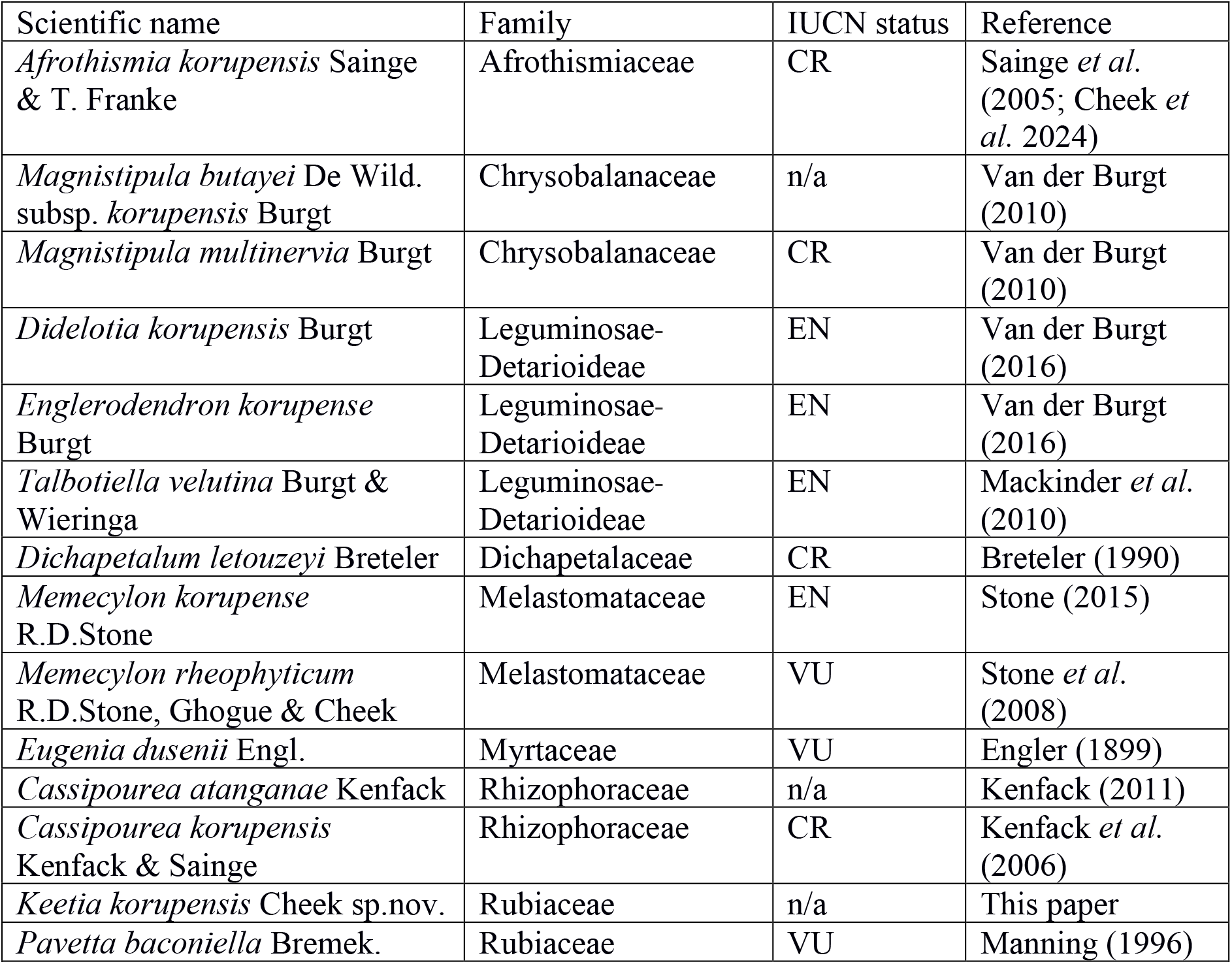
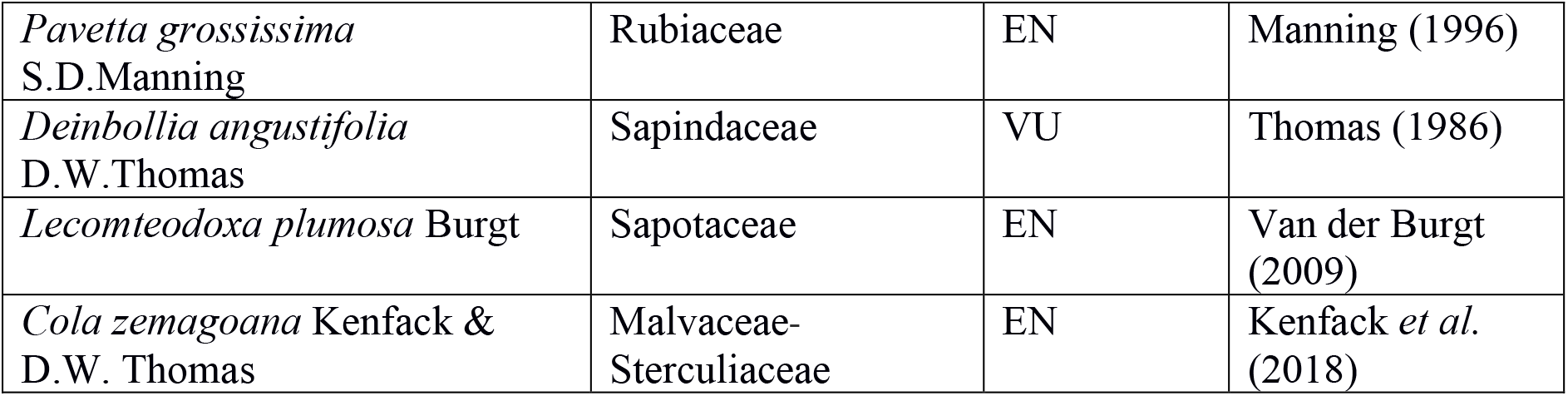
The globally endemic plant species of Korup National Park and their global IUCN extinction risk assessments (from iucnredlist.org, n/a = not available). Data mainly from Murphy *et al*. (2023). Note that *Pavetta baconiella* Bremek. replaces the *nomen nudum P. baconioides* used in error in Murphy *et al*. (2023).

| Scientific name | Family | IUCN status | Reference |
| --- | --- | --- | --- |
| *Afrothismia korupensis* Sainge & T. Franke | Afrothismiaceae | CR | Sainge *et al.* (2005; Cheek *et al.* 2024) |
| *Magnistipula butayei* De Wild. subsp. *korupensis* Burgt | Chrysobalanaceae | n/a | Van der Burgt (2010) |
| *Magnistipula multinervia* Burgt | Chrysobalanaceae | CR | Van der Burgt (2010) |
| *Didelotia korupensis* Burgt | Leguminosae-Detarioideae | EN | Van der Burgt (2016) |
| *Englerodendron korupense* Burgt | Leguminosae-Detarioideae | EN | Van der Burgt (2016) |
| *Talbotiella velutina* Burgt & Wieringa | Leguminosae-Detarioideae | EN | Mackinder *et al.* (2010) |
| *Dichapetalum letouzeyi* Breteler | Dichapetalaceae | CR | Breteler (1990) |
| *Memecylon korupense* R.D.Stone | Melastomataceae | EN | Stone (2015) |
| *Memecylon rheophyticum* R.D.Stone, Ghogue & Cheek | Melastomataceae | VU | Stone *et al.* (2008) |
| *Eugenia dusenii* Engl. | Myrtaceae | VU | Engler (1899) |
| *Cassipourea atanganae* Kenfack | Rhizophoraceae | n/a | Kenfack (2011) |
| *Cassipourea korupensis* Kenfack & Sainge | Rhizophoraceae | CR | Kenfack *et al.* (2006) |
| *Keetia korupensis* Cheek sp.nov. | Rubiaceae | n/a | This paper |
| *Pavetta baconiella* Bremek. | Rubiaceae | VU | Manning (1996) |
| *Pavetta grossissima*<br>S.D.Manning | Rubiaceae | EN | Manning (1996) |
| *Deinbollia angustifolia*<br>D.W.Thomas | Sapindaceae | VU | Thomas (1986) |
| *Lecomteodoxa plumosa* Burgt | Sapotaceae | EN | Van der Burgt (2009) |
| *Cola zemagoana* Kenfack &<br>D.W. Thomas | Malvaceae-Sterculiaceae | EN | Kenfack *et al.* (2018) |

Korup is placed in the extreme SW corner of Cameroon adjacent to the border with Nigeria. It benefits from over 5 m rainfall p.a. in the south, to 1.5 m p.a. in the North. The habitat is largely lowland evergreen rainforest on highly leached impoverished soils, interfacing with Atlantic mangrove in the south and small areas of submontane or cloud forest near the centre where Mt Juahan rises to 1072 m alt. (Murphy *et al*. 2023).

The southern part of the Korup lowland evergreen forest is globally well-known for a series of ecological studies by multiple institutions led by the University of Bern on grove-forming or monodominant species of detarioid legumes (van der Burgt *et al*. 2021). In such areas individual species can form 50% or more of the forest canopy, in patches of various sizes, breaking the standard paradigm of tropical forests. In one area studied, of about 32 km^2^, among the 42 species of detarioid tree recorded, 29 species have been found to be monodominant, a number vastly higher than anywhere else globally (van der Burgt *et al*. 2021). Elsewhere in tropical African forest where such monodominance occurs, typically only a single species is recorded, usually *Gilbertiodendron dewevrei* (de Wild.) J. Léonard (van der Burgt *et al*. 2021). Of the 29 species that are monodominant in Korup, nine species were discovered and published in the course of the ecological research in Korup. Three of those nine species are globally unique to Korup (see Table 2). In addition to these detarioid legume species, new species to science have been discovered and published in the course of inventorying the plots set up to study the monodominant species. These have contributed an additional four endemics to Korup (see the species of Chrysobalanaceae and Sapotaceae in Table 2).

In a parallel programme to the forgoing, led by the Smithsonian Tropical Research Institution, a 50 Ha plot was set up to map and monitor all plant individuals above 1 cm diam. (Thomas *et al*. 2003). This study also contributed the discovery of several new lowland forest species to science including the three Korup endemic trees or treelets of Rhizophoraceae and Sterculiaceae, but also the endemic fully mycotrophic (lacking green tissue) herb *Afrothismia korupensis* (see Table 2).

Both these large plot programmes were instigated by Duncan Thomas who pioneered botanical inventory and conservation in Korup from the early 1980s onwards. His collections from that period form the basis for some of the earliest published Korup endemics such as *Deinbollia angustifolia* (Thomas 1986) as well as the later published *Memecylon rheophyticum* (Stone *et al*. 2008).

An additional network of smaller plots to the north of the forgoing larger plots were studied near the centre of Korup near Mt Juahan, in April 1996 (Cheek & Cable 1998) resulted in the publication of e.g. *Leptonychia moyesiae* Cheek (Cheek *et al*. 2013), and the new genus and species *Korupodendron songweanum* Litt & Cheek (Litt & Cheek 2002) but did not produce any new endemic species for Korup.

Many near endemic species (present in Korup and one or two other locations) brought to light by these studies are generally more highly threatened than those that are endemic to Korup e.g. *Glumea korupensis* Burgt and *Gambeya korupensis* Ewango & Kenfack which both also occur just outside the National Park, (van der Burgt & Newbery 2006; Ewango *et al*. 2016). This is because their sites outside Korup are usually much more highly threatened than those inside Korup. A standard pattern for such near endemics is that their location outside Korup is in the western forests of Mt Cameroon c. 80 km to the SE, in either the Mokoko or Onge forest reserves, e.g. *Tessmannia korupensis* Burgt (van der Burgt 2016). Both Mokoko and Onge forests are also based, at least partly, on highly leached impoverished soils. In the other direction, several threatened species first discovered in these western forests of Mt Cameroon and at first seemingly endemic there, were later found in Korup e.g. *Belonophora ongensis* Cheek & S.E.Dawson (Cheek & Dawson 2000), *Cola suboppositifolia* Cheek (Cheek 2002) and *Drypetes burnleyae* (Cheek *et al*. 2021). However, the western forests of Mt Cameroon also have global endemics that have not been found in the lowland forests of Korup e.g. to endemics to Mokoko such as *Octoknema mokoko* Gosline & Malécot (Gosline & Malécot 2011) and *Dracaena mokoko* Mwachala & Cheek (Mwachala & Cheek 2012). One near endemic, *Gilbertiodendron newberyi* Burgt, is disjunct between Korup and Ebo forest in Littoral Region (van der Burgt *et al*. 2015).

Submontane forest habitat in Korup is confined to Mt Juahan which provided the discoveries of *Newtonia duncanthomasii* Mackinder & Cheek (Mackinder & Cheek 2003) and the likely threatened *Deinbollia oreophila* Cheek (Cheek & Etuge 2009). Both are now known to be widespread in submontane locations outside Korup. These other submontane locations have numerous other threatened submontane species (e.g. Cheek and Csiba (2002) and Stoffelen *et al*. (1997)) which have not been found in Korup, nor have any strictly endemic montane forest species, probably because the area of that habitat in Korup is comparatively small and because only the lower part of the altitudinal range of that habitat is present in Korup.

The earliest known Korup endemic plant species to be recorded are specific to rheophytic habitat (falls and rapids) along the Mana River (also known as the Ndian) that forms part of the National Park’s eastern border in the southern part of the park and has numerous rapids and falls. These records date from the German colonial period more than a century ago, when early botanical collectors such as Dusen and Mildbraed presumably walked up the river after landing by boat at the coast, discovering *Eugenia dusenii* and *Pavetta baconiella*. More recent discoveries of Korup rheophytic endemics are *Deinbollia angustifolia* and *Memecylon rheophyticum*. Together, the rheophytic community of Korup contributes a significant proportion (>20%) of the total number of Korup endemics (4/18) and in terms of woody species is probably the most species diverse rheophytic site in Africa. Cameroon also holds the most species diverse site in Africa for the specialist (obligate) herbaceous rheophytic family Podostemaceae. This is the Lobé falls S of Kribi in S Region (Cheek *et al*. 2017).

Podostemaceae are also represented in Korup’s Mana River with e.g. the highly range restricted near endemic and Endangered *Pohliella laciniata* Engl. (formerly treated as *Saxicolella laciniata* (Engl.) C. Cusset (Cheek *et al*. 2022)).

## Conclusion

Cameroon has the highest number of globally extinct plant species of all countries in continental tropical Africa (Humphreys *et al*. 2019). The extinction of Cameroonian endemic species such as *Oxygyne triandra* Schltr. (Thismiaceae, Cheek *et al*. 2018b) and *Afrothismia pachyantha* Schltr. (Afrothismiaceae, Cheek & Williams 1999; Cheek *et al*. 2019; Cheek *et al*. 2024) are well known examples, recently joined by species such as *Vepris bali* Cheek (Rutaceae, Cheek *et al*. 2018c), *Vepris montisbambutensis* Onana (Onana & Chevillotte 2015) and *Ardisia schlechteri* Gilg (Murphy *et al*. 2023). However, another 127 potentially globally extinct Cameroon species have recently been documented (Onana in Murphy *et al*. 2023: 18 – 22).

It is critical now to detect, delimit and formally name species such as *Keetia korupensis* as new to science, since until they are scientifically recognised, they are essentially invisible to science, and only when they have a scientific name can their inclusion on the IUCN Red List be facilitated (Cheek *et al*. 2020). Most (77%) species named as new to science in 2023 are already threatened with extinction (Brown *et al*. 2023). Many new species to science have evaded detection until today because they are in genera that are long overdue full taxonomic revision as is the case with *Keetia*, or because they have minute ranges which have remained unsurveyed until recently.

If further global extinction of plant species is to be avoided, effective conservation prioritization is crucial, backed up by investment in protection of habitat, ideally through reinforcement and support for local communities who often effectively own and manage the areas concerned. Important Plant Areas (IPAs) programmes, often known in the tropics as TIPAs (Couch *et al*. 2019; Darbyshire (continuously updated); *et al*. 2017; *et al*. 2023; Murphy *et al*. 2023) offer the means to prioritize areas for conservation based on the inclusion of highly threatened plant species, among other criteria. Such measures are vital if further species extinctions are to be avoided of highly rare, threatened species such as *Keetia korupensis*.

## Acknowledgements

Diane Bridson and an anonymous reviewer are thanked for constructive comments on an earlier draft of the paper. Xander van der Burgt, Marcella Corcoran and Moses Elangwe are thanked for support during fieldwork by LP. All authors thank Yvette Berenice Harvey for her searches on their behalf.

The authors declare no conflict of interest

## References

Beentje, H. & Cheek, M. (2003). Glossary. In: Beentje, H. (ed), Flora of Tropical East Africa. Balkema, Lisse.

Breteler, F. J. (1990). Une nouvelle espèce de Dichapetalum Thouars (Dichapetalaceae) du Cameroun. Bull. Mus. Natl. Hist. Nat., B, Adansonia Sér. 4, 11(4): 333

Bridson, D. M. (1985). The reinstatement of Psydrax (Rubiaceae, subfam. Cinchonoideae tribe Vanguerieae) and a revision of the African species. Kew Bull. 40: 687–725. 10.2307/4109853

Bridson, D. M. (1986). The Reinstatement of the African genus Keetia (Rubiaceae, Cinchonoideae, Vanguerieae). Kew Bull. 41(4): 956–994. 10.2307/4102996

Bridson D.M. (1994). A new species of Keetia (Rubiaceae-Vanguerieae) Kew Bull. 49: 803–807. 10.2307/4118075

Bridson D.M. (1998). Rubiaceae (Tribe Vanguerieae) Flora Zambesiaca 5(2): 1–377. <10.2307/4111186 >

Bridson, D. M. & Robbrecht, E. (1993). A spiny-fruited new Keetia (Rubiaceae, Vanguerieae) from Kivu (Zaire). Belg. Journ. Bot. 126: 29–32.

Bridson, D. M. & Verdcourt, B. (1991). Flora of Tropical East Africa–Rubiaceae, 3. Rotterdam/Brookfield, A.A.Balkema. 10.1201/9780203755860

Brown, M., Bachman, S., & Lughadha, E. N. (2023). Three in four undescribed plant species are threatened with extinction. New Phytologist. 10.1111/nph.19214

Cable, S. & Cheek, M. (1998). The Plants of Mt Cameroon, a Conservation Checklist. Royal Botanic Gardens, Kew.

Chapman, J. & Chapman, H. (2001). The Forests of Taraba and Adamawa States, Nigeria an Ecological Account and Plant Species Checklist. University of Canterbury: Christchurch, New Zealand. pp. 221.

Cheek, M. (2002) Three new species of Cola (Sterculiaceae) from western Cameroon, Cameroon. Kew Bull. 57: 402–415. 10.2307/4111117

Cheek, M. (2006). A New Species of Keetia (Rubiaceae-Vanguerieae) from Western Cameroon. Kew Bull. 61(4): 591–594.

Cheek, M. & Bissiengou, P. (2024) Keetia gordonii sp. nov. (Rubiaceae -Vanguerieae) a new species of threatened forest liana from the littoral forest of Gabon. bioRxiv 10.1101/2024.04.18.590134

Cheek M., Bridson D.M. (2019). Three new threatened Keetia species (Rubiaceae), from the forests of the Eastern Arc Mts, Tanzania. Gardens Bulletin Singapore 71(Suppl.2): 155–169. 10.26492/gbs71(suppl.2).2019-12

Cheek, M. & Cable, S. (1997). Plant Inventory for conservation management: the Kew-Earthwatch programme in Western Cameroon, 1993–96, pp. 29–38 in Doolan, S. (Ed.) African Rainforests and the Conservation of Biodiversity, Earthwatch Europe, Oxford.

Cheek, M., & Cable, S. (1998). Preliminary results of the botanical inventory of the Ekundu Kundu region of the Korup National Park. In Proceedings of Workshop on Research and Conservation in Korup National Park and Project Area. Korup Project, Douala, sCameroon (pp. 72–80).

Cheek, M. & Csiba, L. (2002). A new epiphytic species of Impatiens (Balsaminaceae) from western Cameroon. Kew Bull. 57: 669–674. 10.2307/4110997

Cheek, M., & Dawson, S. (2000). A Synoptic Revision of Belonophora (Rubiaceae). Kew Bull. 55(1), 63–80. 10.2307/4117761

Cheek, M. & Etuge, M. (2009). A new submontane species of Deinbollia (Sapindaceae) from Western Cameroon and adjoining Nigeria. Kew Bull. 64: 503–508. 10.1007/s12225-009-9132-4

Cheek, M. & Onana, J.M. (2024). Keetia nodulosa sp. nov. (Rubiaceae-Vanguerieae) of West-Central Africa: bacterial leaf nodulation discovered in a fourth genus and tribe of Rubiaceae. Webbia. Journal of Plant Taxonomy and Geography 79(1): 31–46. 10.36253/jopt-15946

Cheek, M., Achoundong, G., Onana, J-M., Pollard, B., Gosline, G., Moat, J., Harvey, Y.B. (2006). Conservation of the Plant Diversity of Western Cameroon. In: Ghazanfar SA, H.J. Beentje (eds). Proceedings of the 17th AETFAT Congress, Addis Ababa. Ethiopia, 779–791.

Cheek, M., Arcate, J., Choung, H.S. et al. (2013). Three new or resurrected species of Leptonychia (Sterculiaceae-Byttneriaceae-Malvaceae) from West-Central Africa. Kew Bull. 68: 579–590 10.1007/s12225-013-9469-6

Cheek, M., S. Cable, F.N. Hepper, N. Ndam & J. Watts. (1996). Mapping plant biodiversity on Mt. Cameroon. pp. 110–120 in van der Maesen, van der Burgt & van Medenbach de Rooy (Eds), The Biodiversity of African Plants (Proceedings XIV AETFAT Congress). Kluwer. 10.1007/978-94-009-0285-5_16

Cheek, M., Darbyshire, I. & Onana, J.M. (2023a). Discovery and conservation of Monanthotaxis bali (Annonaceae) a new Critically Endangered (possibly extinct) montane forest treelet from Bali Ngemba, North West Region, Cameroon. Kew Bull. 78: 259–270 10.1007/s12225-023-10117-9

Cheek, M., Edwards, S. & Onana, J.M. (2023c). A massive Critically Endangered cloud forest tree, Microcos rumpi (Grewiaceae) new to science from the Rumpi Hills, SW Region, Cameroon. Kew Bull 78: 247–258. 10.1007/s12225-023-10119-7

Cheek, M., Etuge, M. & Williams, S. (2019). Afrothismia kupensis sp. nov. (Thismiaceae), Critically Endangered, with observations on its pollination and notes on the endemics of Mt Kupe, Cameroon. Blumea 64: 158–164. 10.3767/blumea.2019.64.02.06

Cheek, M., Feika, A., Lebbie, A., Goyder, D., Tchiengue, B., Sene, O., Tchouto, P., van der Burgt, X. (2017). A synoptic revision of Inversodicraea (Podostemaceae). Blumea 62: 125–156. 10.3767/blumea.2017.62.02.07

Cheek, M., Gosline, G. & Onana, J.M. (2018c). Vepris bali (Rutaceae), a new critically endangered (possibly extinct) cloud forest tree species from Bali Ngemba, Cameroon. Willdenowia 48: 285–292. 10.3372/wi.48.48207

Cheek, M., Harvey, Y.B., Onana, J-M. (2010). The Plants of Dom. Bamenda Highlands, Cameroon: A Conservation Checklist. Royal Botanic Gardens, Kew.

Cheek M, Harvey Y, Onana J-M. (2011). The Plants of Mefou Proposed National Park. Yaoundé, Cameroon: A Conservation Checklist. Royal Botanic Gardens, Kew.

Cheek, M., Magassouba, S., Molmou, D., Doré, T.S., Couch, C., Yasuda, S., Gore, C., Guest, A., Grall, A., Larridon, I., Bousquet, I.H., Ganatra, B., Gosline, G. (2018a). A key to the species of Keetia (Rubiaceae - Vanguerieae) in West Africa, with three new, threatened species from Guinea and Ivory Coast. Kew Bull. 73: 56. 10.1007/s12225-018-9783-0

Cheek, M., Molmou, D., Magassouba, S., Ghogue, J.-P. (2022). Taxonomic Monograph of Saxicolella (Podostemaceae), African waterfall plants highly threatened by Hydro-Electric projects, with five new species. Kew Bull.,1–31. 10.1007/s12225-022-10019-2

Cheek, M., Ndam, N., & Budden, A. (2021). Notes on the threatened lowland forests of Mt Cameroon and their endemics including Drypetes burnleyae sp. nov., with a key to species of Drypetes sect. Stipulares (Putranjivaceae). Kew Bull. 1–12. 10.1007/s12225-021-09947-2

Cheek, M., Nic Lughadha, E., Kirk, P., Lindon, H., Carretero, J., Looney, B.,Douglas, B., Haelewaters, D., Gaya, E., Llewellyn, T., Ainsworth, M.,Gafforov, Y., Hyde, K., Crous, P., Hughes, M., Walker, B.E., Forzza, R.C., Wong, K.M., Niskanen, T. (2020). New scientific discoveries: plants and fungi. Plants, People Planet 2: 371–388. 10.1002/ppp3.10148

Cheek, M., Onana, J-M., Pollard, B.J. (2000). The Plants of Mount Oku and the Ijim Ridge, Cameroon, a Conservation Checklist. Royal Botanic Gardens, Kew.

Cheek, M., Osborne, J., van der Burgt, X. et al. (2023b). Impatiens banen and Impatiens etugei (Balsaminaceae), new threatened species from lowland of the Cross-Sanaga Interval, Cameroon. Kew Bull. 78: 67–82 10.1007/s12225-022-10073-w

Cheek, M., Pollard, B.J., Darbyshire, I, Onana, J.M. & Wild, C. (2004). The Plants of Kupe, Mwanenguba and the Bakossi Mts, Cameroon. A Conservation Checklist. Royal Botanic Gardens, Kew.

Cheek, M., Sidwell, K., Sunderland T. & Faruk, A. (1992). A Botanical Inventory of the Mabeta-Moliwe Forest. Royal Botanic Gardens, Kew; report to Govt. Cameroon from O.D.A.

Cheek, M., Soto Gomez, M., Graham, S. W., & Rudall, P. J. (2024). Afrothismiaceae (Dioscoreales), a new fully mycoheterotrophic family endemic to tropical Africa. Kew Bull. 79(1): 55–73. 10.1007/s12225-023-10124-w

Cheek, M., Tsukaya, H., Rudall, P.J., Suetsugu, K. (2018b). Taxonomic monograph of Oxygyne (Thismiaceae), rare achlorophyllous mycoheterotrophs with strongly disjunct distribution. PeerJ 6: e4828. 10.7717/peerj.4828

Couch, C., Cheek, M., Haba, P., Molmou, D., Williams, J., Magassouba, S., Doumbouya, S. and Diallo, M.Y., (2019). Threatened habitats and tropical important plant areas (TIPAs) of Guinea, West Africa. Kew: Royal Botanic Gardens, Kew. https://kew.iro.bl.uk/concern/books/ce6950c8-5ed7-4115-b6d4-c09a45b686ff?locale=en

Couch, C., Molmou, D., Magassouba, S., Doumbouya, S., Diawara, M., Diallo, M.Y., Keita, S.M., Koné, F., Diallo, M.C., Kourouma, S. and Diallo, M.B. (2022). Piloting development of species conservation action plans in Guinea. Oryx, pp.1–10. 10.1017/s0030605322000138

Darbyshire, I. (continuously updated) Tropical Important Plant Areas. http://science.kew.org/strategic-output/tropical-important-plant-areas

Darbyshire, I., Anderson, S., Asatryan, A., Byfield, A., Cheek, M., Clubbe, C., Ghrabi, Z., Harris, T., Heatubun, C. D., Kalema, J., Magassouba, S., McCarthy, B., Milliken, W., Montmollin, B. de, Nic Lughadha, E., Onana, J.M., Saidou, D., Sarbu, A., Shrestha, K. & Radford, E. A. (2017). Important Plant Areas: revised selection criteria for a global approach to plant conservation. Biodivers. Conserv. 26: 1767–1800. 10.1007/s10531-017-1336-6.

Darbyshire, I., Kordofani, M., Farag, I., Candiga, R. and Pickering, H. (2015). The Plants of Sudan and South Sudan. Royal Botanic Gardens, Kew.

Darbyshire, I., Richards, S., Osborne, J., Matimele, H., Langa, C., Datizua, C., Massingue, A., Rokni, S., Williams, J., Alves, T. and De Sousa, C., (2023). Important Plant Areas of Mozambique. Kew: Royal Botanic Gardens. https://kew.iro.bl.uk/concern/books/c60f1a8b-07b5-4a7a-9e7f-211b48586faf?locale=zh

Davies, N.M.J., Drinkell, C, Utteridge, T.M.A. (2023). The Herbarium Handbook. Kew Publishing

Engler, A. (1899). Myrtaceae in Diagnosen neuer Africanischer Pflanzen. Notizbl. Königl. Bot. Gart. Berlin 2: 288–292

Ewango, C. E., Kenfack, D., Sainge, M. N., Thomas, D. W., & van der Burgt, X. M. (2016). Gambeya korupensis (Sapotaceae: Chrysophylloideae), a new rain forest tree species from the Southwest Region in Cameroon. Kew Bull. 71: 1–6.

Gosline, G. & Malécot, V. (2011). A monograph of Octoknema (Octoknemaceae—Olacaceae sl). Kew Bull. 66: 367–404. 10.1007/s12225-011-9293-9

Gosline, G., Bidault, E., van der Burgt, X. et al. (2023a). xA Taxonomically-verified and Vouchered Checklist of the Vascular Plants of the Republic of Guinea. Sci. Data 10: 327 (2023). 10.1038/s41597-023-02236-6

Gosline, G. et al. (2023b). Checklist of the Vascular Plants of the Republic of Guinea– printable format (1.10). Zenodo. 10.5281/zenodo.7734985

Gosline, G., Cheek, M., Onana, J.M., Ngansop, Tchatchouang, E., van der Burgt, X.M., MacKinnon, L., Dagallier, L.M.J. (2022). Uvariopsis dicaprio (Annonaceae) a new tree species with notes on its pollination biology, and the Critically Endangered narrowly endemic plant species of the Ebo Forest, Cameroon. PeerJ. Jan 6;9:e12614. 10.7717/peerj.12614

Harvey, Y.B., Pollard, B.J., Darbyshire, I., Onana, J.-M., Cheek, M. (2004). The Plants of Bali Ngemba Forest Reserve. Cameroon: A Conservation Checklist. Royal Botanic Gardens, Kew.

Harvey, Y.B., Tchiengue, B., Cheek, M. (2010). The Plants of the Lebialem Highlands, a Conservation Checklist. Royal Botanic Gardens, Kew.

Humphreys, A.M., Govaerts, R., Ficinski, S.Z., Lughadha, E.N. and Vorontsova, M.S. (2019). Global dataset shows geography and life form predict modern plant extinction and rediscovery. Nature Ecology & Evolution 3.7: 1043–1047. 10.1038/s41559-019-0906-2

IPNI (continuously updated). The International Plant Names Index. http://ipni.org/ (accessed: 07/2023).

IUCN. (2012). IUCN Red List Categories and Criteria: Version 3.1. Second edition.–Gland, Switzerland and Cambridge, UK: IUCN. Available from: http://www.iucnredlist.org/ (accessed: April 2024).

Janssens, S.B., Taedoumg, H., Dessein, S. (2022) Impatiens smetsiana, another example of convergent evolution of flower morphology in Impatiens. Plant Ecology and Evolution 155(2): 248–260. 10.5091/plecevo.89701

Jongkind, C. H. & Lachenaud, O. (2022). Novelties in African Apocynaceae. Candollea 77(1): 17–51. 10.15553/c2022v771a3

Jongkind, C. H. (2002). Two New Species of Keetia (Rubiaceae) from West Africa. Kew Bull. 57(4): 989–992. 10.2307/4115730

Kenfack, D. (2011). Cassipourea atanganae sp. nov., a new species of Rhizophoraceae from Lower Guinea. Adansonia 33(2): 209–213.

Kenfack, D., Sainge, M. N., & Thomas, D. W. (2006). A new species of Cassipourea (Rhizophoraceae) from western Cameroon. Novon: A Journal for Botanical Nomenclature 16(1): 61–64.

Kenfack, D., Sainge MN, Chuyong GB, Thomas DW (2018) The genus Cola (Malvaceae) in Cameroon’s Korup National Park, with two novelties. Plant Ecology and Evolution 151(2): 241–251. 10.5091/plecevo.2018.1410

Lachenaud, O. Luke, Q. Bytebier B. (2017). Keetia namoyae (Rubiaceae, Vanguerieae), a new species from eastern Democratic Republic of Congo. Candollea 72: 23–26. 10.15553/c2017v721a2

Lachenaud, O., Paiva, J., Covelo, F., Cheek, M., & Onana, J. M. (2024). Voacanga madureirae (Apocynaceae), a new species from Atlantic Central Africa. Kew Bull., 1–7. 10.1007/s12225-024-10179-3

Lantz, H. & Bremer, B. (2004). Phylogeny inferred from morphology and DNA data: characterizing well-supported groups in Vanguerieae (Rubiaceae). Botanical Journal of the Linnean Society 146: 257–283. 10.1111/j.1095-8339.2004.00338.x

Litt, A. & Cheek, M. (2002). Korupodendron songweanum, a new genus of Vochysiaceae from West-Central Africa. Brittonia 54(1): 13–17. 10.1663/0007-196x(2002)054[0013:ksanga]2.0.co;2

Mackinder, B. & Cheek, M. (2003). A new species of Newtonia (Leguminosae-Mimosoideae) from Cameroon. Kew Bull. 58: 447–452. 10.2307/4120627

Mackinder, B. A., Wieringa, J. J., & van der Burgt, X. M. (2010). A revision of the genus Talbotiella Baker f. (Caesalpinioideae: Leguminosae). Kew Bull. 65: 401–420. 10.1007/s12225-010-9217-0

Maisels, F.M., Cheek, M., Wild, C. (2000). Rare plants on Mt Oku summit, Cameroon. Oryx 34: 136–140. 10.1017/s0030605300031057.

Manning, S. D. (1996). Revision of Pavetta subgenus Baconia (Rubiaceae: Ixoroideae) in Cameroon. Annals of the Missouri Botanical Garden, 87–150. 10.2307/2399970

Murphy, B., Onana, J.M., van der Burgt, X. M., Tchatchouang Ngansop, E., Williams, J., Tchiengué, B., Cheek, M. (2023). Important Plant Areas of Cameroon. Royal Botanic Gardens, Kew.

Mwachala, G. & Cheek, M. (2012) Dracaena mokoko sp. nov. (Dracaenaceae-Ruscaceae/Asparagaceae) a Critically Endangered forest species from Mokoko, Cameroon. Nordic J. Bot. 30(4): 389–393. 10.1111/j.1756-1051.2011.01487.x

Onana, J.-M. (2011). The Vascular Plants of Cameroon. A Taxonomic Checklist with IUCN Assessments. Flore Du Cameroun 39. Ministry of Scientific Research and Innovation, Yaoundé, Cameroon.

Onana, J.M. & Cheek, M. (2011). Red Data Book of the Flowering Plants of Cameroon, IUCN Global Assessments. Royal Botanic Gardens, Kew.

Onana, J.M. & Chevillotte, H. (2015). Taxonomie des Rutaceae–Toddalieae du Cameroun revisitée: découverte de quatre espèces nouvelles, validation d’une combinaison nouvelle et véritable identité de deux autres espèces de Vepris Comm. ex A. Juss. Adansonia sér. 3. 37: 103–129. https://doi.org/c10.5252/a2015n1a7

POWO (continuously updated). Plants of the World Online. Facilitated by the Royal Botanic Gardens, Kew. http://www.plantsoftheworldonline.org (downloaded 1 Dec. 2023).

Quintanar, A., Sonké, B., Simo-Droissart, M., Barberá, P., Libalah, M., Harris, D.J. (2023). A matter of warts: a taxonomic treatment for Drypetes verrucosa (Putranjivaceae, Malpighiales) and a new cauliflorous species from Cameroon and Nigeria, D. stevartii. Plant Ecology and Evolution 156(2): 160–173. 10.5091/plecevo.102004

Sainge, M. N., Franke, T. & Agerer, R. (2005). A new species of Afrothismia (Burmanniaceae, tribe Thismieae) from Korup National Park, Cameroon. Willdenowia 35: 287–291. doi:10.3372/wi.35.35209

Shang, H., & Zhang, L. B. (2023). Systematics of the Fern Genus Didymochlaena (Didymochlaenaceae). Systematic Botany 48(1): 110–139. 10.1600/036364423X16758873924144

Sosef, M.S.M., Wieringa, J.J., Jongkind, C.C.H., Achoundong, G., Azizet Issembé, Y., Bedigian, D., Van Den Berg, R.G., Breteler, F.J., Cheek, M., Degreef, J. (2006). Check-list des plantes vasculaires du Gabon. Scripta Botanica Belgica 35. National Botanic Garden of Belgium. 435 pp.

Stoffelen, P., Cheek, M., Bridson, D., Robbrecht, E. (1997). A new species of Coffea (Rubiaceae) and notes on Mt Kupe (Cameroon). Kew Bull. 52: 989–994. 10.2307/4117826

Stone, R. D. (2015). Taxonomic treatment of Memecylon L. section Felixiocylon RD Stone (Melastomataceae), with descriptions of four new species from Cameroon, Gabon, and Equatorial Guinea (Bioko). Adansonia 37(1), 47–61.

Stone, R. D., Ghogue J.-P. & Cheek, M. (2008). Revised treatment of Memecylon sect. Afzeliana (Melastomataceae-Olisbeoideae), including three new species from Cameroon. Kew Bull. 63: 227–241. 10.1007/s12225-008-9033-y

Stone, R.D., Tchiengué, B., Cheek, M. (2023). The endemic plant species of Ebo Forest, Littoral Region, Cameroon with a new Critically Endangered cloud forest shrub, Memecylon ebo (Melastomataceae-Olisbeoideae). bioRxiv 12.20.572583 10.1101/2023.12.20.572583

Tabi, E. A., Younchahou, M. N. & Mohanty, A. (2020). The effect of armed conflict on biodiversity and its implication on wildlife: a case study on the Lebialem Highlands, South-West Region, Cameroon. Int. J. Sci. Res. Biol. Sci. 7: 107–112. https://www.isroset.org/pdf_paper_view.php?paper_id=2190&13-ISROSET-IJSRBS-05008.pdf

Thiers, B. (continuously updated). Index Herbariorum: A global directory of public herbaria and associated staff. New York Botanical Garden’s Virtual Herbarium. [continuously updated]. – Available from: http://sweetgum.nybg.org/ih/ (accessed: Feb. 2023).

Thomas, D. W. (1986). Notes on Deinbollia species from Cameroon. Annals of the Missouri Botanical Garden 73(1): 219–221.

Thomas, D.W., Kenfack D., Chuyong G.B., et al. (2003). Tree species of Southwestern Cameroon: tree distribution maps, diameter tables, and species documentation of the 50-hectare Korup Forest Dynamics Plot. Smithsonian Tropical Research Institute, Washington, D.C. Available from https://forestgeo.si.edu/sites/default/files/korup.pdf

Turland, N.J., Wiersema, J.H., Barrie, F. R., Greuter, W., Hawksworth, D.L., Herendeen, P.S., Knapp, S., Kusber, W.-H., Li, D.-Z., Marhold, K., May, T.W., McNeill, J., Monro, A.M., Prado, J., Price, M.J. & Smith, G.F. (ed.) (2018). International Code of Nomenclature for algae, fungi, and plants (Shenzhen Code) adopted by the Nineteenth International Botanical Congress Shenzhen, China, July 2017. – Glashütten: Koeltz Botanical Books. [= Regnum Veg. 159]

van der Burgt, X.M. (2009). Lecomtedoxa plumosa (Sapotaceae), a new tree species from Korup National Park, Cameroon. Kew Bull. 64: 313–317 10.1007/s12225-009-9103-9

van der Burgt, X.M. (2010). Two new taxa in Magnistipula (Chrysobalanaceae) from Korup National Park, Cameroon. Plant Ecology and Evolution 143(2): 191–198. 10.5091/plecevo.2010.400

van der Burgt, X. M. (2016). Didelotia korupensis and Tessmannia korupensis (Leguminosae, Caesalpinioideae), two new tree species from Korup National Park in Cameroon. Blumea-Biodiversity, Evolution and Biogeography of Plants 61(1): 51–58. doi:10.3767/000651916X691402

van der Burgt, X.M. & Newbery D.M. (2006). Gluema korupensis (Sapotaceae), a new tree species from Korup National Park, Cameroon. Kew Bull. 61(1): 79–84. https://www.jstor.org/stable/20443247

van der Burgt, X.M., Mackinder B.A., Wieringa J.J. & Estrella M. de la (2015). The Gilbertiodendron ogoouense species complex (Leguminosae–Caesalpinioideae), Central Africa. Kew Bull. 70: 29. 10.1007/s12225-015-9579-4

van der Burgt, X. M., Newbery, D. M., & Njibili, S. (2021). The structure of Leguminosae-Detarioideae dominant rain forest in Korup National Park, Cameroon. Plant Ecology and Evolution 154(3): 376–390. doi: 10.5091/plecevo.2021.1879

